# *In vivo* Effects of Temperature on the Heart and Pyloric Rhythms in the Crab, *Cancer borealis*

**DOI:** 10.1101/249631

**Authors:** Dahlia Kushinsky, Ekaterina O. Morozova, Eve Marder

**Affiliations:** Biology Department and Volen Center Brandeis University Waltham, MA 02454

**Keywords:** photoplethysmography, crustaceans, stomatogastric nervous system, cardiac ganglion, central pattern generators, Q_10_

## Abstract

**Summary Statement:** Temperature elevation increases the frequency of the heart and pyloric rhythms of the crab, *Cancer borealis*, but the heart rhythm has a higher critical temperature than the pyloric rhythm.

**Abstract:** The heart and pyloric rhythms of crustaceans have been studied separately and extensively over many years. Local and hormonal neuromodulation and sensory inputs onto these central pattern generating circuits play a significant role in the animals’ responses to perturbations, but are usually lost or removed during *in vitro* studies. To examine simultaneously the *in vivo* motor output of the heart and pyloric rhythms, we used photoplethysmography (PPG). In the population measured (n = 49), the heart rhythm frequencies ranged from 0.3–2.3 Hz. The pyloric rhythms varied from 0.2–1.6 Hz. During multiple hour-long recordings, many animals held at control temperature showed strong inhibitory bouts in which the heart decreased in frequency or become quiescent and the pyloric rhythm also decreased in frequency. Many animals show significant coherence in frequency between the rhythms at the frequency of the heart rhythm. We measured the simultaneous responses of the rhythms to temperature ramps by heating or cooling the saline bath while recording both the heart and pyloric muscle movements. Q_10_s, critical temperatures (temperatures at which function is compromised), and changes in frequency were calculated for each of the rhythms tested. The heart rhythm was more robust to high temperature than the pyloric rhythm.

## Introduction

Central pattern generators (CPGs) are responsible for the generation of many rhythmic motor patterns that are vital for muscle movements of an animal’s life. In crustaceans, mechanisms of central pattern generation have been studied using the cardiac ganglion (CG), which produces heart movements, and the stomatogastric ganglion (STG) which generates the rhythmic movements of the stomach (Cooke, 2002; Maynard, 1972). These two rhythms are subject to both local and global modulation. These rhythms have been studied separately, both *in vivo* and in dissected and isolated preparations, but now we ask whether these rhythms appear to be coordinated in an intact animal.

The heart of the crustacean is neurogenic, controlled by motor neuron discharges produced by the CG. In normal conditions, the heart continuously pumps hemolymph through an open circulatory system to distribute oxygen and neuromodulators to the animal’s tissues (McMahon, 1995; McMahon, 2001a; McMahon, 2001b). This circulatory system is formed by elaborate capillary beds in many of the animal’s tissues, paired with valves that allow for selective hemolymph distribution (McMahon, 1995; McMahon, 2001a; McMahon, 2001b). The cardiac output, calculated as the stroke volume/heart rate, must be maintained to ensure adequate hemolymph distribution (McGaw and McMahon, 1996). Hemolymph flow rate and stroke volume are subject to wide variations both within and across animals (McGaw and McMahon, 1996). Changes in flow rate are likely due to modifications in stroke volume or valve activation, and may occur in response to local tissue demands (McGaw and McMahon, 1996; Wilkens and McMahon, 1994). Changes in cardiac output may also be the result of spontaneous firing events in the CNS, CG, or cardioarterial valves of the heart (McGaw and McMahon, 1996; Wilkens and McMahon, 1994).

The CG, controlling the movement of the heart, contains an intrinsically oscillating pacemaker to form a central pattern generated rhythm. Additionally, the heart is innervated by extrinsic fibers, one inhibitory and two excitatory, that modify heart rate in relation to behavioral demands (Cooke, 2002). The CG of the crab, *Cancer borealis*, is comprised of nine neurons, 4 small and 5 large cells, that burst in time to produce muscle movements. The CG also produces patterned bursts of impulses in response to simple stimuli, such as excitation from stretch sensitive dendrites (Cooke, 2002). This allows the heart to adjust its frequency and strength of contractions when faced with different metabolic needs (Dickinson et al., 2016a; Dickinson et al., 2015a).

The stomatogastric nervous system (STNS) has long been used to study central pattern generation and mechanisms underlying rhythmic and continuous motor patterns (Maynard, 1972). The STNS controls the movement of the crustacean stomach (Maynard and Dando, 1974; Morris and Hooper, 1997; Morris and Hooper, 1998; Morris and Hooper, 2001). Foregut movements allow for feeding behavior, including chewing, swallowing, and processing of waste (Clemens et al., 1998b; Johnson and Hooper, 1992). Oxygen tension alters the neuronal activity of the STG and therefore its motor output (Clemens et al., 2001). The stomach is a complex mechanical structure with ossicles that provide mechanical support and insertions for intrinsic stomach muscles (Maynard and Dando, 1974). The STNS generates the continuously active and rapid pyloric rhythm and the slower, episodic gastric mill movements (Clemens et al., 1998a; Clemens et al., 1999; Clemens et al., 1998b; Clemens et al., 2001; Clemens et al., 1998c; Heinzel, 1988; Heinzel and Selverston, 1988; Rezer and Moulins, 1983; Soofi et al., 2014).

The pericardial organs (POs) are secretary organs that deliver neurohormones to the animal’s hemolymph (Alexandrowicz and Carlisle, 1953; Chen et al., 2010; Christie et al., 1995; DeKeyser et al., 2007; Hui et al., 2012). These neuromodulators may increase the frequency and amplitude of the heartbeat (Christie et al., 2008; Cruz-Bermudez and Marder, 2007; Sullivan and Miller, 1984; Williams et al., 2013), or change the frequency of the pyloric rhythm (Hooper and Marder, 1987; Marder, 2012; Marder and Bucher, 2007). Hemolymph containing neuromodulators flows into the heart and is pumped throughout the open circulatory system, filling the ophthalmic artery, directly bathing the STG. The modulators released into the hemolymph act hormonally, and therefore may affect the animal’s various organs in the same manner, or may have varied effects depending on the receptors present in the tissues. This provides a feedback system within the animal, allowing for the simultaneous adjustment of organ activity when needed in the animal.

Throughout its life, an organism must respond to perturbations that affect the nervous system and the biomechanical structures it controls. Crustaceans are poikilotherms and therefore may experience wide variations in body temperature due to temperature changes in the environment. Because of this, it is relevant to study the effect of temperature on the activity of the STNS and CG, as temperature changes may simultaneously affect cellular processes of both systems and disrupt neuronal function. Throughout their lives, crustaceans experience both short term temperature fluctuations, due to changing tidal patterns during a single day, and long term temperature fluctuations, due to seasonal temperature variations (Soofi et al., 2014; Tang et al., 2010; Tang et al., 2012). Despite these varied temperature changes, animals must maintain rhythmicity and performance of the foregut and heart. Therefore, temperature is a useful manipulation to study network stability in the face of perturbation.

Previous work on crustaceans indicate that both the heart and pyloric rhythms are robust to temperature changes. In the heart, studies have shown that the strength of a heartbeat decreases and heart rate increases with increases in temperature (Camacho et al., 2006; Worden et al., 2006). Therefore, the increase in heart rate partially compensates for the decrease in stroke volume as the CG is further pushed from its normal state at 11°C. The pyloric triphasic rhythm is maintained across a wide range of temperatures. In both the heart and the pyloric rhythm, the maximum frequency attained at highest temperatures *in vivo* are lower than those of *in vitro* conditions, likely due to the presence of sensory feedback and neurohormonal input (Soofi et al., 2014; Worden et al., 2006).

The Q_10_ is a measure of the sensitivity of a biological process to a 10°C change in temperature. Many biological processes have Q_10_s between two and three, while some temperature sensitive ion channels have Q_10_s as high as 50 or 100 (Marder et al., 2015). If all Q_10_s of the components involved in a biological process are similar, this process is likely to be temperature compensated (Caplan et al., 2014; Marder et al., 2015; O’Leary and Marder, 2016; Robertson and Money, 2012). In both the heart and pyloric rhythms, the Q_10_s for the rhythm frequencies are similar *in vivo* and *in vitro* (Soofi et al., 2014; Worden et al., 2006). Worden and colleagues (2006) noted that in the heart, the Q_10_s of various parameters were 1-3.5. The pyloric rhythm increases in frequency with Q_10_s between 2 and 2.5 while conserving its phase relationships within the triphasic rhythm (Soofi et al., 2014; Tang et al., 2010).

Temperature increases beyond 23°C lead to severely disrupted motor patterns and subsequent loss of rhythms in the heart and pyloric rhythms (Rinberg et al., 2013; Tang et al., 2012). Recordings of the pyloric rhythm show that preparations appear similar at permissive temperatures, but extreme temperatures disrupt, or crash them, in dissimilar ways revealing their variability (Marder et al., 2015; Rinberg et al., 2013; Tang et al., 2012).

In this study we systematically explore the potential relationships between stomach and heart rhythms, and ask whether they are coordinately sensitive to perturbation by temperature.

## Materials and Methods

### Animals

Adult male Jonah crabs *(Cancer borealis)* between 400 and 700 grams were obtained from Commercial Lobster (Boston, MA). Animals were housed in tanks with artificial seawater (Instant Ocean) between 10°C and 13°C on a 12 hour light/dark cycle without food for a maximum of 10 days. For *in vivo* experiments, animals were placed in a 25 liter tank filled with approximately 10 liters of artificial seawater inside an incubator at 10°C to 12°C. Experiments were done between 9/1/16 and 7/12/17.

Prior to each experiment, crabs were weighed and anesthetized on ice for 10 minutes. Photoplesmogram (PPG) sensors (Vishay CNY70331) (Fig. 1), as described in Depledge (1983), were placed on the carapace above the heart and pyloric muscles to record the heart and pyloric rhythms, respectively. Sensors were secured to the carapace using dental wax and cyanoacrylate glue (Starbond, EM-2000) and covered in Marine Adhesive Sealant (3M, Fast Cure 5200) to waterproof and ensure the stability of the sensors over time. After the sensors were attached to the animal, we waited a minimum of 16 hours prior to experimental recordings, during which time the animals were not handled and the door to the incubator was not opened.

**Figure 1.**
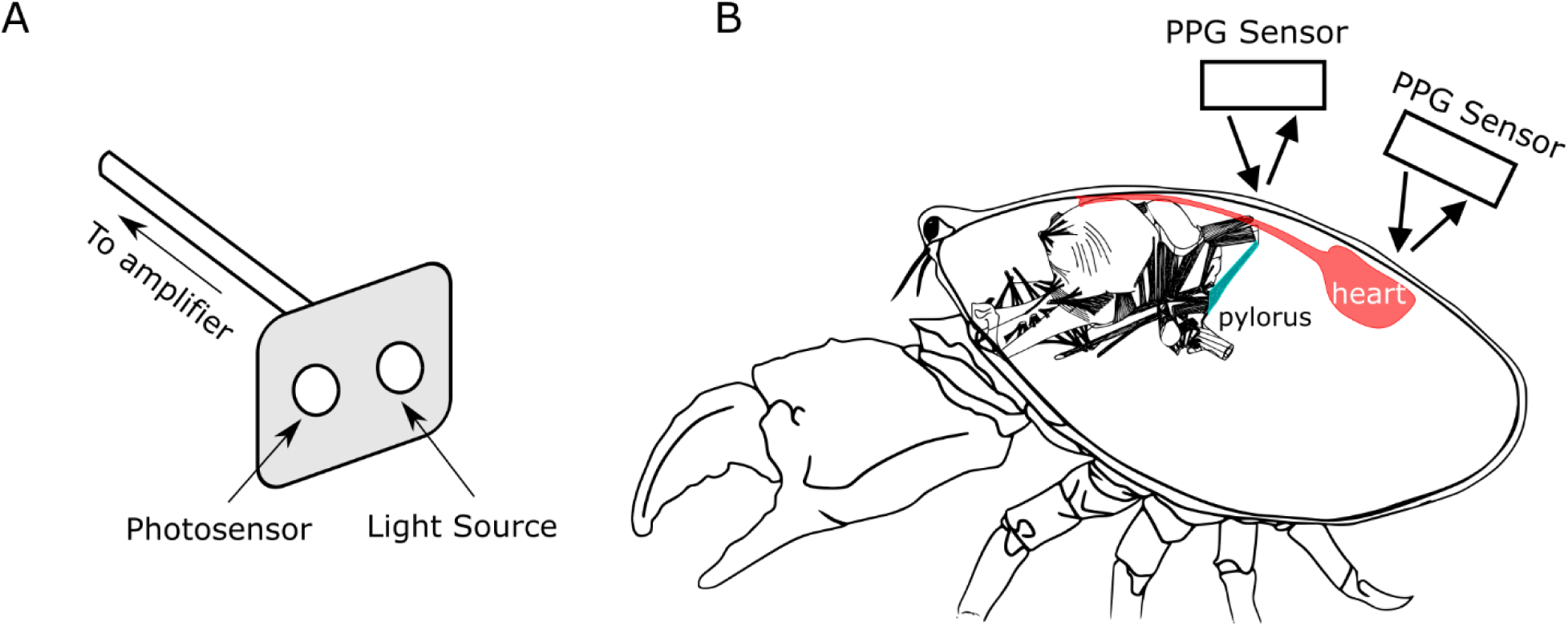
Use of photoplesmography (PPG) allows for continuous noninvasive recordings of the heart and pyloric muscles. **A**) Drawing of PPG sensor system, indicating light source and photosensor on device. **B**) Drawing of the PPG sensor positioning for detection of heart and pylorus movements when placed on the carapace of an animal. Infrared light from the sensors is emitted and travels through the animal’s carapace before being reflected from the muscle directly below the PPG sensor. Crab image modified from http://stg.rutgers.edu/Resources.html. Stomach image was modified from Maynard and Dando, (1974)·

### Temperature Experiments

After a period of baseline (10°C to 12°C) recording, water temperature was manipulated by flowing either cold or warm saline into the tank through a tube inserted through the door of the incubator. A vacuum line was used to pump water out of the tank to maintain a constant volume. Temperature was slowly ramped from 11°C to 32°C over 1.5 to 2 hours. Heart rate was closely monitored to ensure health during the temperature changes and ramps were halted once the heart rate developed an arrhythmia or decreased to baseline frequencies, indicating that a ‘critical temperature’ had been reached. Increasing temperatures past this critical temperature lead to death of the animal, as the heart no longer recovered functionality.

### Data Acquisition and Analysis

PPG data were acquired through the PPG amplifier (Newshift AMP03) and recorded digitally through a digitizer (Axon Digidata 1550B) into computer software (AxoScope 10.6) with a sampling frequency of 500 Hz. Data were analyzed using custom written C and MATLAB codes.

### Analysis of heart rhythm frequency

Heart rhythm frequency was calculated as the frequency at the peak spectral power. We used the Burg (1967) method to estimate the power spectrum density at each moving-time window. The Burg method (1967) fits the autoregressive (AR) model of a specified order p in the time series by minimizing the sum of squares of the residuals. The fast-Fourier transform (FFT) spectrum is estimated using the previously calculated AR coefficients. This method is characterized by higher resolution in the frequency domain than traditional FFT spectral analysis, especially for a relative short time window (Buttkus, 2000). We used the following parameters for the spectral estimation: data window of 12.8 s (128 samples), 50% overlap to calculate spectrogram, number of estimated AR-coefficients p=window/4+1. Before the analysis, the voltage offsets of the PPG recordings were removed, low pass filtered to 5 Hz using six-order Butterworth filter and downsampled. Average baseline frequencies of heart rhythm were calculated as median frequencies at peak of the power spectral density for each window of a spectrogram during baseline conditions. Cumulative histograms of baseline frequencies were calculated as a sum of histograms from individual animals normalized so that the sum of bar heights is less than or equal to 1.

### Analysis of the pyloric rhythm frequency

The pyloric rhythm frequency was calculated in a similar way as the heart frequency in those cases that showed no interference from heart activity. However, there were instances in which the heart activity was influencing the pyloric rhythm. This was obvious in the pyloric rhythm spectrogram as a peak in the power spectrum density at the frequency of heart rhythm. To identify the intrinsic frequency of the pyloric rhythm in cases with heart interference, a linear regression model was fit to the pyloric signal taking into account the phase difference between the heart and pyloric signals. The heart signal was multiplied by the coefficients of the regression model and subtracted from the pyloric signal. Then spectrogram of the subtracted signal was calculated, and the pyloric frequency was identified as the frequency at peak spectral power. In some cases, the pyloric rhythm frequency could not be determined due to irregularities in the signal.

### Analysis of the inhibitory bouts

We used a hidden Markov model (HMM) to infer the active and inhibitory states of heart rhythms. In HMM, a timeseries is modeled as being generated probabilistically from an underlying discrete-valued stochastic process (Rabiner, 1989). The data can be either discrete- or continuous-valued, while the unobservable ‘hidden’ state is a discrete random variable that can take *n* possible values (in our case n=2, representing active and inhibitory states). Estimation of the transition probabilities for HMM was done using the Baum-Welch algorithm, which utilizes an expectation maximization (EM) algorithm (Bilmes, 1998). The initial parameters used for the detection: transition matrix P_AI_ = P_IA_ = 0.9, P_AA_ = P_II_ = 0.1, where P_AI_ is transition probability from active to inhibitory state, P_AI_ is a transition probability from inhibitory to active state, P_ii_ is a transition probability from inhibitory to inhibitory state and P_AA_ is a transition probability from active to active state. Identification by HMM states was used to calculate the durations of active and inhibitory bouts of heart rhythm.

### Coherence analysis

Coherence between heart and pyloric data was calculated with multi-taper Fourier analysis (Mitra et al., 1999) using Chronux toolbox (http://www.chronux.org). Data were binned into two-minute bins, moved in 5 second steps. The time-bandwidth product was set to 10, and 19 tapers were used. Peak coherence and frequency of peak coherence were calculated for each window. The theoretical confidence level of the coherence was calculated as following:

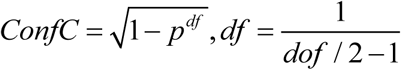
where dof is degrees of freedom. Based on these calculations, coherence values > 0.625 reached the significance of p < 0.05 with a Bonferroni correction. Data sets that had significant coherence more 50% of time during baseline were considered to have significantly coherent heart and pyloric rhythms and were calculated for each data set. The phase difference between the heart and pyloric rhythms was calculated at the frequency of the peak coherence for each window and median value of phase difference was reported for each dataset with significantly coherent signals.

### Q_10_ estimation

We estimated the Q_10_ of frequency of the heart and pyloric rhythms *in vivo*. Frequency (Fr) of the heart and pyloric rhythms were plotted as a function of temperature (T) in a logarithmic scale, and the Q_10_ was extracted from the slope of the linear regression (m) following the equation:

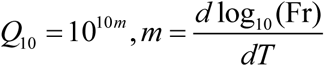

The goodness of fit of the linear regression model for each dataset was assessed by calculating the coefficient of determination R^2^, calculated as R^2^ = (correlation coefficient)^2^. We report R^2^ for heart and pyloric data in the tables below. For majority of the fits we obtained high values of R^2^ > 0.8.

**Table 1.** Statistics for heart frequency dependence on temperature.

| $R^2$ for heart frequency vs T fit | $R^2$ for pyloric frequency vs T fit |
| --- | --- |
| 0.921 |  |
| 0.773 |  |
| 0.792 |  |
| 0.589 | 0.956 |
| 0.486 | 0.332 |
| 0.9052 | 0.8536 |
| 0.9085 | 0.933 |
| 0.9618 | 0.789 |
| 0.98 | 0.9871 |
| 0.996 |  |
| 0.81 | 0.8691 |

### Critical Temperature Analysis

The critical temperature was defined as the temperature at which the heart and pyloric movements became irregular and the frequency of muscle contraction significantly dropped. This was determined from the spectrograms of heart and pyloric rhythms. In some cases, the critical temperature of the pyloric rhythm was impossible to determine due to irregularity in pyloric rhythm signal.

### Statistical analysis

All statistical analyses were done in Matlab. Between group comparisons were done using one-way ANOVA. Significance level was set to 0.05. When appropriate Bonferroni corrections were implemented.

## Results

Heart and pyloric muscle movements of *Cancer borealis* were recorded *in vivo* using photoplesmography (PPG) (Fig. 1). In most of the experiments described, both rhythms were recorded from the same animals, although in some only the heart rhythms were recorded. It is straightforward to place a PPG sensor over the heart because the heart is dorsal, and situated just under the carapace, and its movements are large and vigorous, almost in the same plane as the sensor itself. The dorsal artery exits the heart and travels anteriorly close to the surface of the stomach. The pylorus is the most posterior region of the stomach, but most of its movements are in the interior of the animal. The thin straps of the dorsal dilator muscles of the pyloric region connect the pylorus to insertions just below the carapace, and the movements of these muscles are almost orthogonal to the carapace. Therefore, picking up these movements is considerably more difficult than those of the heart. Each heart beat would lead to a pressure wave in the artery, and the artery runs close to the dorsal dilator muscles, so it is to be expected that the pyloric PPG sensor might pick up some trace of the heart rhythm.

Figure 2 shows sample recordings and spectrograms calculated from them for both the heart and pyloric rhythms. Figure 2A (left) shows three short stretches of raw data from 3 crabs; animal 1 had a relatively slow and steady heart rhythm (0.6 Hz), animal 2 had a relatively high frequency (1.1 Hz) and steady heart rhythm, and animal 3 showed a rhythm that switched from high to low frequency spontaneously, and then from low to high frequency in response to a saline injection. To the right of each of the sets of raw traces there is a spectrogram showing 1–2 hours of recordings, with the location of each of the raw data indicated.

**Figure 2.**
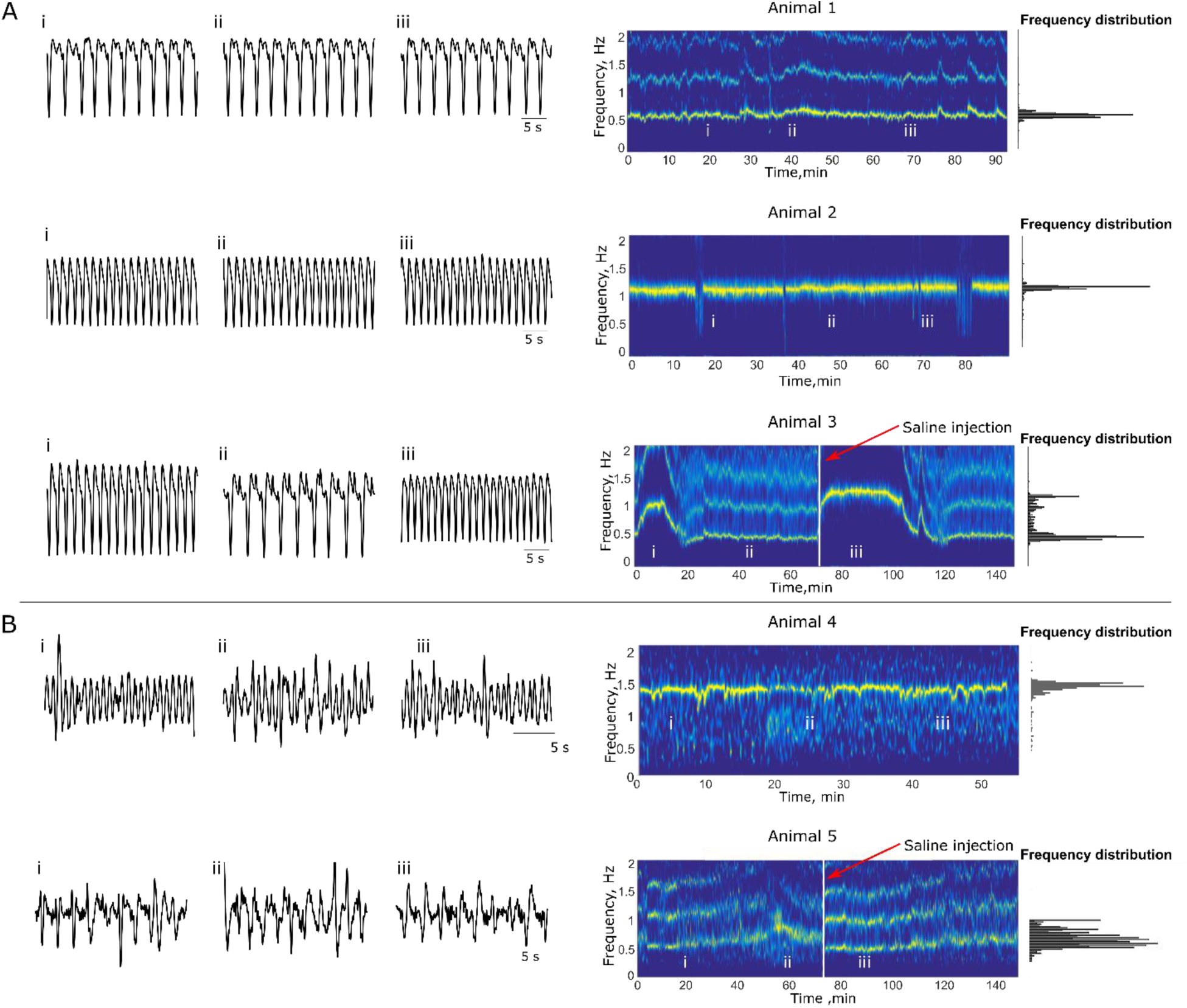
Variability of heart and pyloric rhythm frequency at baseline conditions. **A**) Examples of cardiac rhythm recordings from three animals at different times during baseline conditions (left) with their spectrograms (right). Roman numerals mark where 30-second raw traces were taken from in the longer baseline frequency data (spectrograms). Animals 1 and 2 have stable low and high frequency cardiac rhythms respectively. The heart frequency of animal 3 switches between low and high frequency states spontaneously during baseline and after the saline injection (indicated by vertical white line). **B**) Examples of pyloric rhythm recordings from two animals during baseline conditions (left) with their spectrograms (right). Animal 4 has a stable high frequency pyloric rhythm. Animal 5 has a more variable pyloric rhythm. The pyloric rhythm frequency decreases after the saline injection (indicated by vertical white line). Note that the waveforms of the pyloric rhythms are more complex than the waveforms of the heart rhythm due to the complex movement patterns of the pyloric muscles.

Recordings of the pyloric rhythm with PPGs yielded complex, highly variable waveforms due to the angle of the dorsal dilator muscle movement with respect to the PPG sensor placement on the carapace of the animal. Figure 2B shows raw trace plesmograph recordings and their associated spectrograms of the pyloric rhythms from two animals. Animal 4 maintained a stable frequency of pyloric rhythm, while animal 5 showed a more variable rhythm during its baseline period. The frequency of the pyloric rhythm tends to decrease in response to saline injection as in the spectrogram for animal 5.

Figure 3 summarizes pooled frequency data for the heart from 49 animals and the pylorus from 29 animals. In all cases, the data came from stretches of recordings in excess of 30 minutes. All of the pyloric rhythm data came from animals that were also used for heart measurements. The histogram in Figure 3A shows an apparent multimodal distribution of heart frequencies (Hartigan’s dip test, dip=0.067, p = 0.029). Heart rhythm frequencies ranged between 0.4 Hz and 2.4 Hz (Fig. 3A). No correlation was found between heart rhythm frequency and time of year of experiment (p = 0.25, Pearson’s correlation coefficient). Additionally, no correlation was found between average heart rhythm frequency and weight of crab (p = 0.37, Pearson’s correlation coefficient).

**Figure 3.**
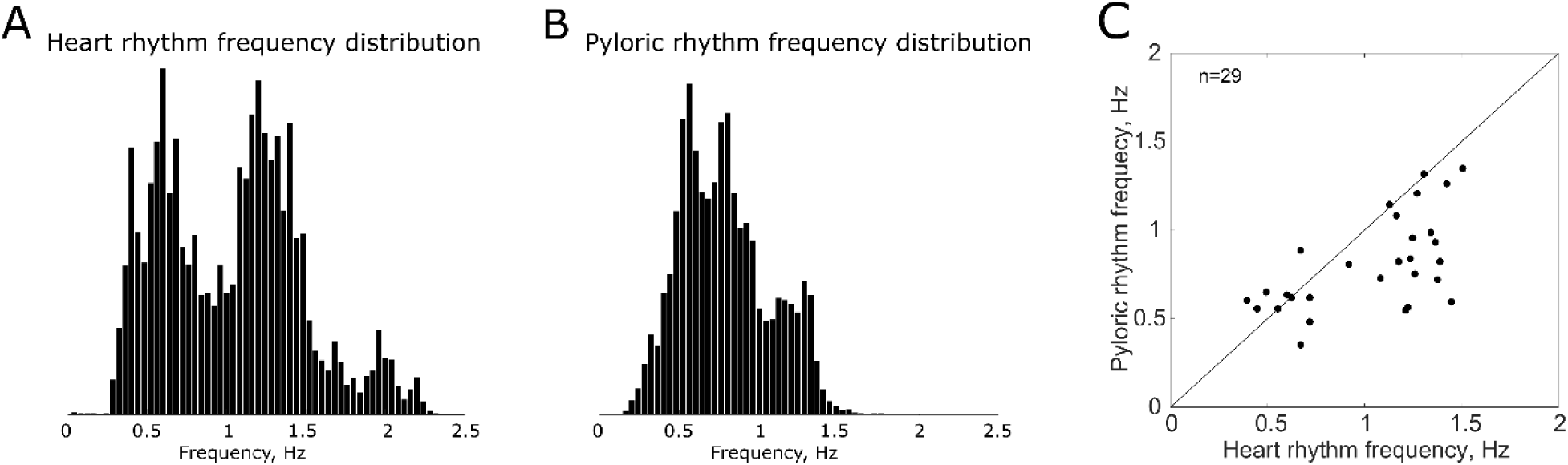
Frequency distributions of the heart and pyloric rhythms. **A**) The sum of normalized frequency distributions of heart rhythms of 49 animals at baseline conditions is multimodal. **B**) Sum of normalized frequency distributions of pyloric rhythms of 29 animals at baseline conditions. The spread in frequencies of the pyloric rhythms is smaller than of heart rhythms. **C**) Plot of baseline heart rhythm frequency versus pyloric rhythm frequency for each animal for which both were recorded. The pyloric frequency on average is lower than the heart rhythm frequency.

Figure 3B shows a more normal distribution of the pyloric rhythm frequencies. The spread in pyloric rhythm frequencies was much smaller than in heart frequencies, ranging from 0.2 Hz to 1.6 Hz. In Figure 3C, we plotted the frequency of the pyloric rhythm as a function of the heart rhythm for the 29 animals for which we had measurements of both. Note that more than half of the points are not found close to the identity line, suggesting that the pyloric and heart recordings are picking up rhythms in the same general frequency range, but are not identical.

### Inhibitory bouts

Heart rhythms often displayed periods of bradycardia, during which the heart considerably slowed or halted for a significant period (Fig. 4). We defined these periods as inhibitory bouts using a hidden Markov model (procedure described in the methods section). Periods of bradycardia were marked by a decrease in both amplitude and frequency (Fig. 4) of heart rhythm PPG recordings by at least 33% that lasted at least 10 seconds. An example of a 24 hr recording with multiple inhibitory bouts can be seen in Figure 4A. Termination of inhibitory bouts was associated with a return of amplitude and frequency. A temporary increase in amplitude of the heart signal could sometimes be observed immediately following the inhibitory bout (Fig A inset). Inhibitory bouts were seen in 20/49 animals of the population tested (41%). In most cases, when inhibitory bouts were seen, they occurred repeatedly over extended periods of time, such as seen in the 24 hr recordings shown in Figure 4. Bout durations and frequency were variable both across and between animals (Fig. 4B). The occurrence of inhibitory bouts was not significantly correlated with the time of year (p = 0.79, Pearson’s correlation coefficient) or weight of animal (p = 0.41, Pearson’s correlation coefficient).

**Figure 4.**
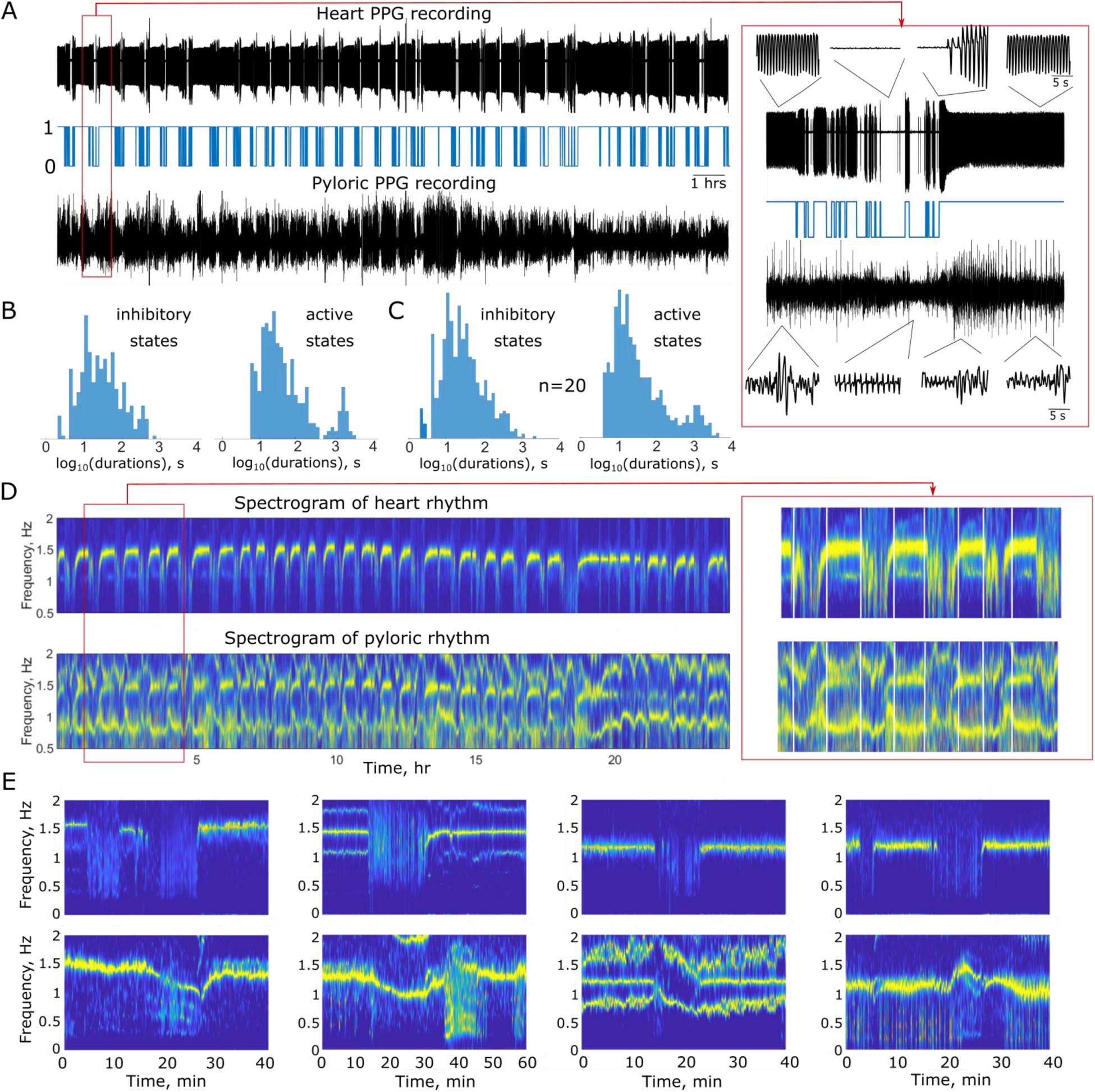
The heart rhythm exhibits inhibitory bouts that influence the pyloric rhythm. **A**) Top panel, example of a 24 hour recording of the heart rhythm showing inhibitory bouts throughout the recording session. Middle panel shows states identified from the heart rhythm trace using hidden Markov model. State 1 corresponds to an active state and state 0 corresponds to an inhibitory bout. Bottom panel, simultaneously recorded pyloric rhythm. Expanded traces during single inhibitory bout are shown in the red box. **B**) Distribution of durations of active and inhibitory states of heart rhythm shown in A. **C**) Cumulative distribution of durations of active and inhibitory states of heart activity of 20 animals. Heart rhythms of 20 out of 49 animals displayed inhibitory bouts with a mean duration of 30 s. Active state durations display bimodal distribution. **D**) Top panel shows spectrogram of the heart rhythm shown in A. The heart rhythm frequency in between the inhibitory bouts is maintained relatively constant at 1.5 Hz. Bottom panel shows spectrogram of the pyloric rhythm shown in A. The pyloric rhythm decreases in frequency during the heart inhibitory bouts followed by the increase in frequency at the end of the bouts. Zoomed in portion of the spectrogram is shown in the red box. **E**) More examples of the frequency changes of the pyloric rhythm during and following the heart inhibitory bouts from four different animals. Top panels are the spectrograms of heart activity and bottom panels are the spectrograms of the pyloric activity. These data feature interaction between the cardiac and the pyloric activity on the time scale of minutes.

Long periods of bradycardia affected the frequency and amplitude of pyloric rhythm. Simultaneously recorded pyloric and heart signals in long-term experiments show that the amplitude of the pyloric signal decreases during the inhibitory bout of the heart (Fig 4A). Spectral analysis revealed that the frequency of the pyloric rhythm modestly decreases during the heart inhibitory bouts (Fig. 4 C). However, there is considerable variability in the changes in frequency of pyloric rhythm, as can be seen from the spectrograms calculated during single inhibitory heart bout in 4 animals (Fig 4 D). For example, the experiments shown in the first panel illustrates an example when the pyloric rhythm was reliably moving with a frequency of 1.5 Hz while the heart was beating at 1.6 Hz. In this animal when the heart temporarily stopped, the pyloric rhythm slowed to 0.9 Hz. The second example also shows a strong decrease in pyloric frequency during the heart inhibitory bout. The third example showed a transient increase followed by a decrease, while the fourth example showed a slight increase in frequency.

<u>Relationships between the heart and pyloric rhythms</u>. Although it is clear that the pyloric rhythm and heart rhythms are often at different frequencies, and the pyloric rhythm continues during the inhibitory bouts, spectrograms of the pyloric rhythm frequently reveal a band at the heart frequency. This is very clearly illustrated in the third example in Fig. 4D, where the heart rhythm is seen as a tight band at about 1.2 Hz. That same band is seen below in the pyloric rhythm traces. When the heart stops, the heart band disappears from both recordings.

To look at the potential influence of the heart rhythm on the pyloric rhythm, we calculated the time-frequency coherence between simultaneously recorded heart and pyloric rhythms in 2-minute bins moved in 5 second steps. In 70% of the animals, the heart and pyloric rhythms were significantly coherent at the frequency of the heart more than 50% of the time during the baseline period. Because the pyloric frequency often changes when the heart stops, this suggests that some kind of biomechanical coupling or common drive is influencing the two structures. Figure 5 A illustrates an example of such coupling in an individual animal. The coherence peaks at 1.1 Hz frequency (Fig. 5 A), which is the frequency of heart oscillations shown in the spectrogram of the heart signal (Fig 5A). The spectrogram of the pyloric rhythm has two frequency bands: one at the frequency of approximately 0.5 Hz, which is intrinsic frequency of the pyloric oscillations, and another on the heart rhythm frequency. By calculating the phase at the frequency of peak coherence we determined that the pyloric rhythm is shifted by 190 degrees relative to the heart rhythm in this animal. This can also be seen from the cross-correlation function, which has a minimum at 0.075 s lag. We also calculated auto-correlations for the heart and pyloric rhythms. The pyloric rhythm has a more complex auto-correlation function than the heart rhythm featuring two peaks, one peak on the lag corresponding to the period of the heart oscillations and another peak on the period of intrinsic pyloric oscillations. Figure 5 B illustrates one of the few examples when the heart and pyloric rhythms were not coherent. The cross-correlation function of the two rhythms in this example is flat. Overall, 28 out of 40 animals had coherent rhythms with a magnitude of coherence more than 0.6 and phase differences between minus and plus 45 degrees (Fig. 5 C). There was no dependence of the percent of time the rhythms were significantly coherent on the amplitude and frequency of the heart signal (Fig. 5C).

**Figure 5.**
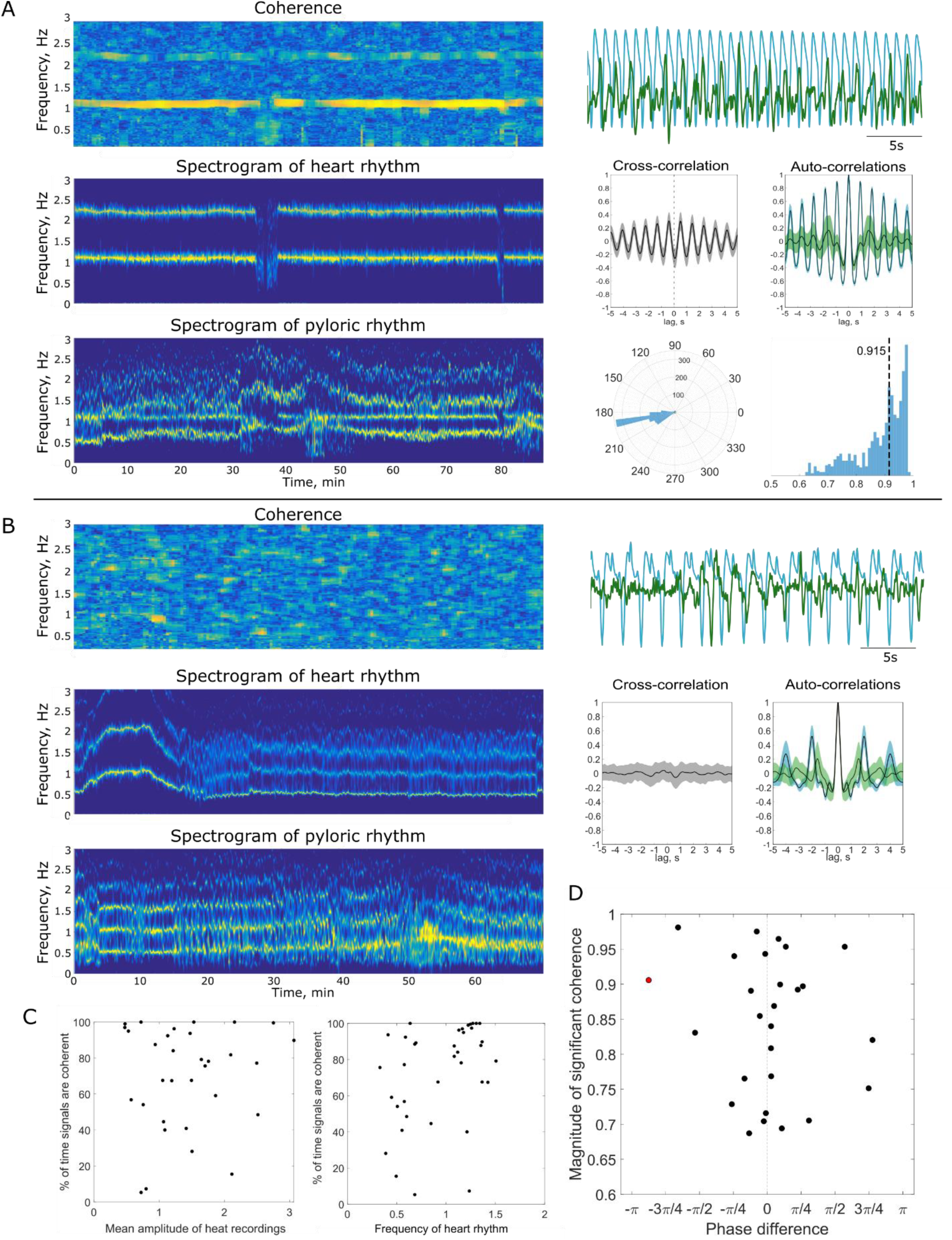
Pyloric and heart rhythms are coherent at the frequency of the heart rhythm in the majority of animals. **A**) An example of high coherence between heart and pyloric rhythms from an individual animal. Top panel shows time-frequency coherence of the heart and pyloric rhythms at baseline. Middle panels show spectrograms of the heart and pyloric rhythms at baseline. Traces on the right show heart rhythm (blue) and pyloric rhythm (green), simultaneously recorded at baseline conditions. Data shown here are 30s segments of the full data range for which spectra and coherence were calculated. Below the traces, heart-pyloric cross-correlation (gray) and autocorrelation functions of heart (blue) and pyloric (green) rhythms are shown. The panel below shows the distribution of phase differences between the heart and pyloric rhythms calculated at 2 minute windows, moved in 10 s steps. The mean phase shift in this dataset is 190 degrees. Last panel, the distribution of magnitudes of peak coherence calculated at 2 minute windows, moved in 10 s steps. Median magnitude of peak coherence is 0.915. **B**) Example of the absence of coherence between the heart and pyloric rhythms from an individual animal. Top panel shows time-frequency coherence of heart and pyloric rhythms at baseline. Middle panels show spectrograms of heart and pyloric rhythms at baseline. Traces on the right are heart rhythm (blue) and simultaneously recorded pyloric rhythm (green) recorded at baseline. Data shown here are 30 s segments of the full data range for which spectra and coherence were calculated. Below the traces heart-pyloric cross-correlation (gray) and autocorrelation functions of heart (blue) and pyloric (green) rhythms are shown. **C**) Coherence statistics from all animals with simultaneous recordings of heart and pyloric rhythms (n=40). The left panel shows the percentage of time pyloric and heart rhythms are significantly coherent at baseline versus the amplitude of the recorded heart signal. The middle panel shows percentage of time pyloric and heart rhythms are significantly coherent at baseline versus the frequency of heart rhythm. **D**) Scatter plot showing median magnitudes of peak coherence versus median phase difference between heart and pyloric rhythms. The pyloric and heart rhythms were significantly coherent >50% of time in 70% of animals.

<u>Effects of temperature on heart and pyloric rhythms</u>. We tested the effects of increasing temperature on the heart movements of twelve animals and on pyloric movements in nine animals (Fig. 6). Figure 6A shows raw data from a typical experiment showing heart and pyloric movements during a temperature ramp from 11°C to 28°C and then back to 11°C. Figure 6B shows plots of the heart and pyloric rhythm frequency for three animals as a function of the temperature. Frequency was calculated for both heart and pyloric rhythms as the frequency at the peak of the power spectrum density of the rhythms in each sliding window. In all the heart movement recordings, the frequency increased with increasing temperature until crashes of the heart rhythm occurred. Rhythm crashes were characterized by the occurrence of low frequency and irregular patterns of activity. Crash of the heart rhythm is followed by the recovery of activity as the temperature decreases. Note that despite the common pattern in which hearts of different animals respond to changes to the temperature, there is across animal variability in the heart rhythm activity in response to almost identical temperature ramps. Particularly, maximal frequencies, critical temperatures and recovery patterns are different in different animals (Fig. 6B). In the pyloric recordings, waveforms changed in overall shape, often exhibiting peaks followed by long plateaued waveforms. Similar to the heart activity, the frequency of the pyloric rhythm also increased with increasing temperature, again characterized by irregular periods of “crashed” activity at high temperatures.

**Figure 6.**
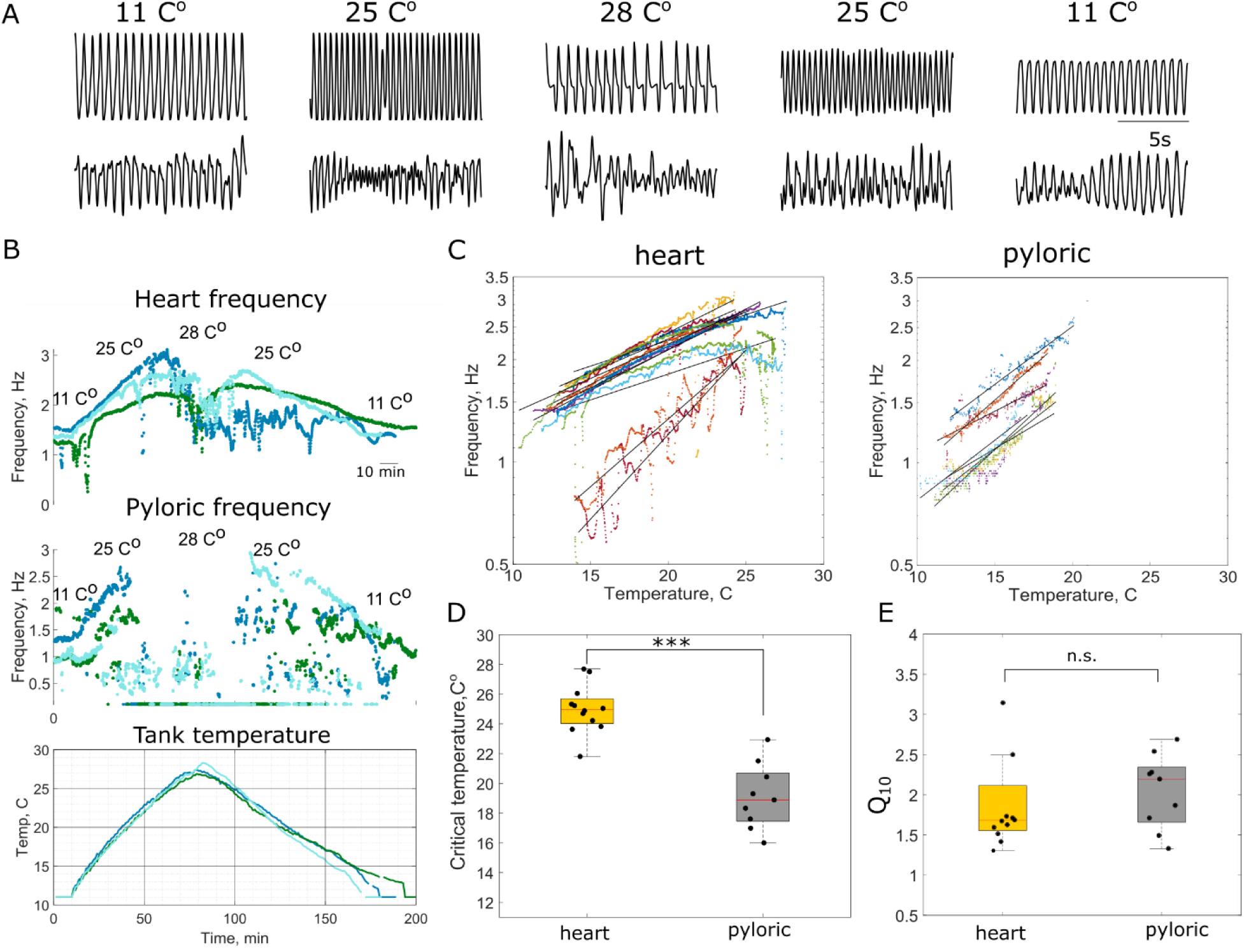
The pyloric rhythm is more sensitive to increases in temperature than is the heart rhythm. **A**) Raw traces of heart and pyloric muscle activity at baseline (11°C), during increasing (25°C), critical (28°C) and decreasing portions (25°C) of the temperature ramp. **B**) Top trace shows the change in frequency of heart rhythms of three animals in response to almost identical temperature ramps shown in the bottom panel. Middle trace shows the change in frequency of simultaneously recorded pyloric rhythm in response to the temperature ramp. The pyloric rhythm is less robust to the increases in temperature than the heart rhythm and crashes at a much lower temperature. “Crash” is evident by significant decrease in frequency and amplitude of the pyloric rhythm. **C**) Frequencies of the heart and pyloric rhythms during increasing portion of temperature ramps plotted as a function of a temperature in a logarithmic scale, each color corresponds to an individual animal. A line was fit to data points for each animal’s heart frequencies to estimate Q_10_s. **D**) Critical temperature of the heart rhythm is significantly higher than of the pyloric rhythm (mean heart critical temperature *T_Hr___critical_ =* 25 ±1.62 *C*^°^, mean pyloric critical temperature *T*_p__*_critical_* = 19.1 ± 2.76, *one-way ANOVA, p=0.0005, F(1,19)=21.07).* **E**) Q_10_s of heart and pyloric frequencies are not significantly different *(mean Q10 of heart frequency is* 2.007 ± 0.854, *mean Q10 of pyloric frequency is* 2.04 ± 0.467, *one-way ANOVA, p=0.9155, F(1,19)=0.01*).

Frequencies of the heart and pyloric rhythms are plotted as a function of temperature in a logarithmic scale in Figure 6C. Linear models were fitted to the data points for each animal to estimate heart and pyloric frequency Q_10_s. The pyloric rhythms consistently crashed at lower temperatures than the heart rhythm in the same animal. The critical temperature was defined as the temperature at which cardiac or pyloric stability and regularity was lost, with a subsequent drop in contraction frequency to near baseline values. Critical temperatures of the heart muscle movements were collected for all twelve animals and for the pyloric muscle movements for nine animals (Fig. 6D). The mean heart critical temperature was 25.0°C (s.d. = 1.62) and the mean pyloric critical temperature was 19.1°C (s.d. = 2.76) (Fig. 6D). The critical temperatures of the heart and pyloric muscle movements were significantly different (*one-way ANOVA, p=0.0005, F(1,19)=21.07)*.

Q_10_, a measurement of the rate of change of a biological process in response to a change in temperature, was calculated for both the heart and pyloric rhythm for each animal tested (Fig. 6E). The frequency of the heart and pyloric rhythms was plotted as a function of environmental temperature in a logarithmic scale and linear regression model was fitted into data points (Fig. 6E). Coefficients of determinations (R^2^) for each fit are shown in table in methods section. Q_10_s were calculated as slopes of linear models. Q_10_s for heart rhythms, ranged from 1.3 to 4.2 with a mean of 2.007 (s.d. = 0.854). Pyloric rhythm Q_10_s ranged from 1.33 to 2.7 with a mean of 2.04 (s.d. = 0.47). The Q_10_s of the heart and pyloric rhythms were not significantly different as determined by *one-way ANOVA (p=0.127, F(1,19)=2.54)*.

Finally, we calculated the coherence between the heart and pyloric rhythms during the temperature ramps to determine whether temperature perturbation affects the relationship between signals. An example time-frequency coherence from an individual animal is shown in Figure 7A. The coherence peaks at the frequency of heart oscillations at baseline as well as during the temperature ramps. Examples of coherence and phase differences calculated at different stages of the experiment (baseline, rising phase of temperature ramp, and decreasing phase of temperature ramp) show that the amplitude of the coherence remains high throughout the whole experiment and the temperature perturbation does not affect the phase relationship between rhythms. In this example the rhythms oscillate in phase (phase difference at the frequency of maximal coherence is shown by arrows in Figure 7C). Peak coherence at baseline and during the rising phase of the temperature ramp as well as the percent of time the rhythms were coherent and their phase difference were calculated for all nine temperature experiments with simultaneously recorded heart and pyloric rhythms (Fig 7D). Coherence between signals was calculated up to the point of the pyloric rhythm crash. There were no statistically significant differences between mean peak coherences at baseline and during the temperature ramps as determined by one-way ANOVA (*p=0.806, F(1,32)=0.06)*. Similarly, there were no statistically significant differences between mean phase differences at baseline and during the temperature ramps as determined by one-way ANOVA *(p=0.759, F(1,32)=0.1)*. Finally, the rhythms were coherent approximately the same amount of time at baseline and during the rising phase of the temperature ramp (one-way ANOVA, *p=0.354, F(1,16)=0.91)*. In the majority of the datasets, the rhythms were coherent 100% of the time. Together, these data suggest that rhythms that are coherent at baseline remain coherent during the increase of environmental temperature without a significant change in phase relationship.

**Figure 7.**
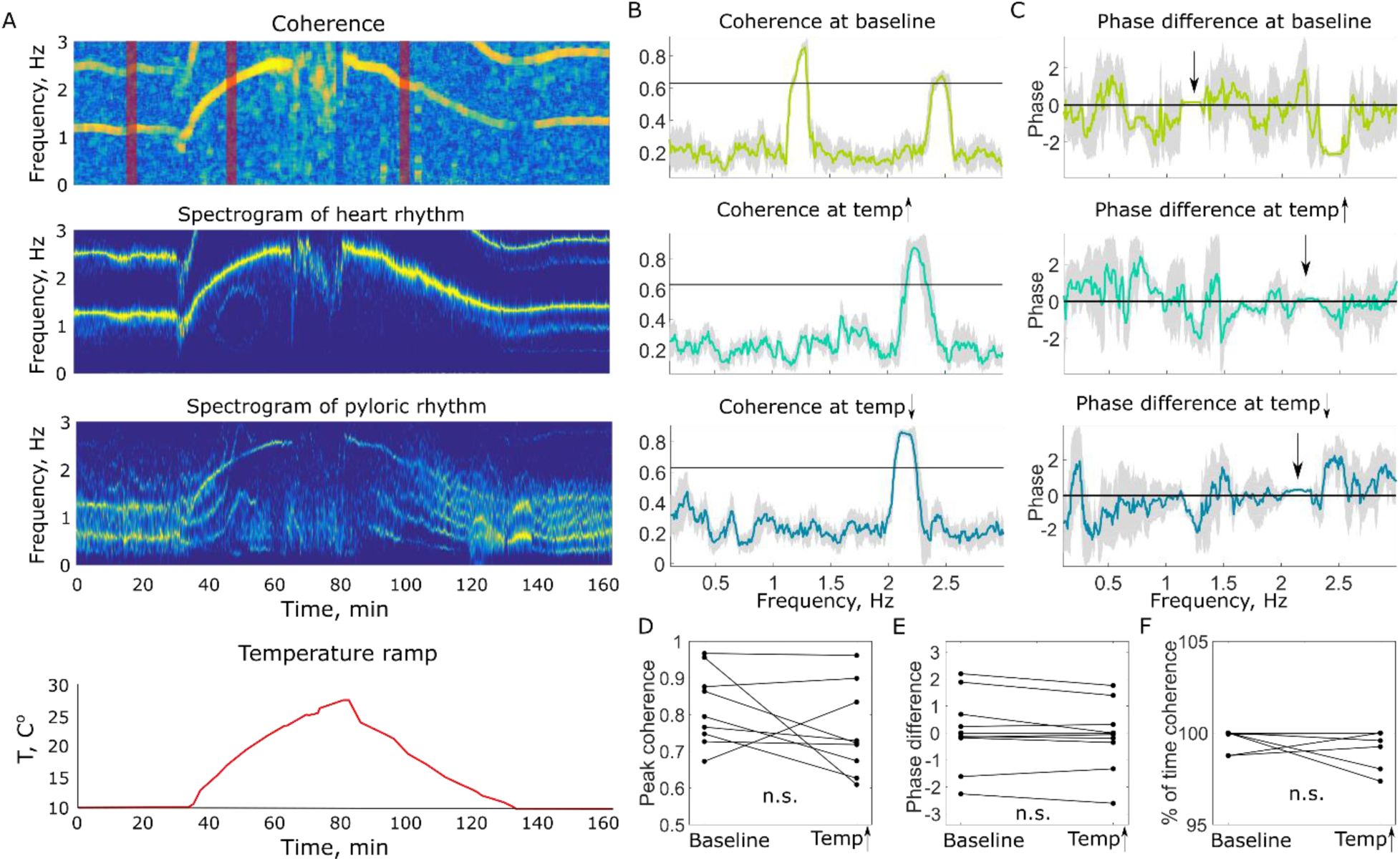
Coherent heart and pyloric rhythms remain coherent during the temperature ramp without change in phase relationship. **A**) An example of coherence between heart and pyloric rhythms and spectrograms of the rhythms at baseline and during temperature ramp from an individual animal. Bottom panel shows temperature in the tank during the experiment. **B**) Examples of coherence at baseline, during rising and decaying phases of temperature ramps. Horizontal black line shows theoretical significance level. The times at which example coherences were taken are shown by the red bars in part A. Rhythms are coherent throughout the whole experiment at frequency of heart rhythm. **C**) Phase difference of the rhythms in A at baseline, during rising and decaying phases of temperature ramps. Phase difference between heart and pyloric rhythm remains constant throughout the whole experiments. **D**) Peak coherence between heart and pyloric rhythms from all temperature experiments (n=9) at baseline and during rising phase of temperature ramp. Coherence during temperature ramp was calculated up to the critical temperature of the pyloric rhythm. There were no statistically significant differences between mean peak coherences at baseline and during the temperature ramp as determined by one-way ANOVA, *p=0.806, F(1,32)=0.06*. **E**) Phase difference between heart and pyloric rhythms were calculated for all temperature experiments (*n*=9). There were no statistically significant differences between mean phase differences at baseline and during temperature ramp as determined by one-way ANOVA, *p=0.759, F(1,32)=0.1.* **F**) Percent of time heart and pyloric rhythms were coherent. There were no statistically significant differences between mean percent of time rhythms were coherent at baseline and during temperature ramp as determined by one-way ANOVA, *p=0.354, F(1,16)=0.91.* In majority of the datasets rhythms were coherent %100 of time.

## Discussion

It is always interesting to determine the extent to which different body rhythms are correlated in their activity. This is especially interesting with respect to the heart and stomach rhythms in crustaceans because the stomatogastric ganglion is situated just anterior to the heart in the dorsal artery, where it is directly perfused by hemolymph containing hormones released into the circulatory system. To the best of our knowledge, ours is the first study to record simultaneously the heart and stomach’s pyloric rhythms both under control conditions and in response to temperature changes.

### Baseline Functioning of the Heart Rhythm

The crustacean heart is neurogenic, and therefore the activity of the cardiac ganglion directly regulates the heart beat frequency. The CG must be able to produce activity regularly and across a wide range of perturbations. It must be modifiable to adjust to the animal’s metabolic needs (Dickinson et al., 2016b; Dickinson et al., 2015b; Robertson and Money, 2012) and reliable enough to ensure that the heart continues to pump hemolymph at all times. Both frequency and contraction amplitude, two factors which must be regulated to ensure the animals’ success, are expressions of the activity of the CG, specifically its interburst interval, rate, and firing patterns.

We captured heart activity using PPG recording techniques while limiting the invasiveness and stress placed on the animal. PPG recordings of the heart musculature were reliable over time and showed differences in frequency of heart beats within the population (Fig. 2). In the majority of experiments, an animal’s baseline heart rate was relatively stable (Fig. 2), but in some animal’s frequency switches were spontaneously seen (Fig. 2A, animal 3). During baseline recordings, animals were not subjected to changes in temperature, light, salinity, stress or environmental factors that could interact with the metabolic needs of the animal or the functions of the CG. During this time, the firing of the CG, and therefore the activity of the heart, would not be expected to appreciably change. Nonetheless, the heart rate of resting animals falls into a multimodal distribution. These data suggest that heart activity, and therefore activity of the CG, falls into states of higher or lower activity. These states may be a consequence of variable metabolic needs of the animal during the time of recording, such as digestion, movement, or excretion. The CG may therefore have mechanisms for switching between high and low activity states through neuromodulation or extrinsic neural input. Alternatively, the high and low activity states may reflect circatidal (rhythms that are governed by the tide) (Chabot and Watson, 2010a) or circadian rhythms the animal experiences (Chabot and Watson, 2010b; De La Iglesia and Hsu Y., 2010).

Several animals displayed heart inhibitory bouts, periods of bradycardia during which the heart slowed or completely stopped for a minimum of 10 seconds. These inhibitory bouts persisted throughout an entire experiment, and likely last through long periods in an animal’s life. Within an animal, inhibitory bouts persist for extended periods of time during a long recording session (Fig. 4B). Inhibitory bouts have previously been observed as linked to times of gill ventilation during reversal of pumping of the scaphognathite (McMahon, 1999). This is likely due to overflow of ocean water, and therefore oxygen, in the gills, causing the organism’s heart to stop for a significant period. It is interesting, however, that this occurs in only about 41% of animals tested. This implies a mechanism affecting both the heart and the gills that influences the overall levels of dissolved O_2_ within the animal.

### Baseline Function of the Pyloric Rhythm

The STG maintains its triphasic rhythm to control the movement of the muscles, and is robust to several global perturbations, including temperature (Soofi et al., 2014; Tang et al., 2010; Tang et al., 2012) and pH (Haley, Hampton and Marder, 2018). Previous work has shown the importance of animal-to-animal variability in responses to such perturbations (Hamood et al., 2015; Hamood and Marder, 2014; Hamood and Marder, 2015). Interestingly, there are differential sensitivities of the responses of isolated STGs and cardiac ganglia to pH, again with the cardiac ganglion being more robust. However, no one has previously compared the *in vivo* responses of the heart and pylorus to any perturbation.

In the experiments presented here, PPG sensors recording the muscle movement of the pylorus were placed on the carapace above the dorsal dilator muscle. While heart rhythm waveforms were relatively simple, with one maximum and minimum, pyloric rhythm waveforms were more complex (Fig. 2B). The complexity of the PPG waveform is likely due to the positioning of the muscles being recorded and their movements in relation to the PPG sensor. *In vivo*, the pyloric rhythm frequency drifts more than that of the heart rhythm frequency over a baseline period. Across the population of tested and analyzed animals, the pyloric rhythm had a more normal distribution, unlike that of the heart rhythm (Fig. 3).

### Comparison of the Heart and Pyloric Rhythms

While the generation and movement of the heart and pylorus have been extensively studied separately in the past, here we examined the potential interactions between these two central pattern generated movements. To determine how these two essential rhythms may interact in a single animal, we calculated time-frequency coherence between the rhythms of animals in controlled environments. In a majority of animals, the heart and pyloric rhythms were coherent at the frequency of the heart rhythm (Fig. 5A). We know that this coherence is not simple cross-talk between the sensors themselves, as the pyloric rhythm continues when the heart stops (Fig. 4C,D), and because there are numerous instances when the two rhythms are quite different. We suspect that the coherence between the two rhythms is a bio-mechanical coupling between the heart and the stomach, which are situated close to each other under the dorsal carapace. Moreover, the dorsal artery runs between the paired dorsal dilator muscles (the source of the pyloric signal) and the PPG sensor placed above the posterior part of the stomach may pick up pulsations of the artery that occur with each heartbeat.

### Temperature as a Perturbation to Expose the Mutual Changes in the Heart and Pyloric Rhythms

Previous work on temperature has shown that both the heart and pyloric rhythms increase in frequency with increases in temperature (Marder et al., 2015; Soofi et al., 2014; Tang et al., 2010; Worden et al., 2006). Both rhythms are also known to crash beyond critical temperatures, at which point the muscles no longer move in canonical patterns (Soofi et al., 2014; Tang et al., 2012), and it is likely that the muscles themselves cannot contract correctly at high temperatures. In intertidal worms, the heart critical temperature likely corresponds with the onset of anaerobic metabolism, due to a mismatch between the animal’s oxygen needs and the supply being delivered to the heart (Zielinski and Portner, 1996).

In the data presented here, it is clear that temperature affects both rhythms and increases their overall frequency, with similar Q_10_s (Fig. 6E). This indicates that increases in temperature cause similar changes in the two frequencies, until a crash point occurs. Interestingly, the heart is more robust to extreme temperature changes than the pyloric rhythm.

Crabs can live for days and weeks without eating, but presumably cannot survive for extended periods of time without hemolymph oxygenation and circulation. From this perspective, it is easy to justify the fact that the critical temperature for the heart rhythm is higher than that for the pyloric rhythm. The additional 4–5°C might make a big difference for an animal caught in shallow water during the summer, and give it time to find its way to more hospitable environments. Interestingly, the mean critical temperatures are within the range that *C. borealis* might experience during a New England summer in shallow water or intertidal adventures. The animal-to-animal variability in these critical temperatures may be important signatures that explain the survival of some animals during summer heat.

## Acknowledgements

We thank Dr. Markus Frederich for an introduction into the use of PPG system, and Jessica Haley for assistance in early data analysis. Data in this paper were first presented for partial fulfilment of requirements for BS/MS program (DK) at Brandeis University.

## References

Alexandrowicz, J. S. and Carlisle, D. B. (1953). Some experiments on the function of the pericardial organs in Crustacea. J. Mar. Biol. Assoc. U.K. 32, 175–192.

Bilmes, J. A. (1998). A Gentle Tutorial of the EM Algorithm and its Application to Parameter Estimation for Gaussian Mixture and Hidden Markov Models. In International Computer Science Institute, pp. 7–13. Berkeley, CA.

Burg, J. P. (1967). Maximum entropy spectral analysis. In Proc. 37th Meeting Society of Exploration Geophysicist. Oklahoma City, OK.

Buttkus, B. (2000). Spectral Analysis and Filter Theory. Berlin: Springer.

Camacho, J., Qadri, S. A., Wang, H. and Worden, M. K. (2006). Temperature acclimation alters cardiac performance in the lobster *Homarus americanus*. J Comp Physiol A Neuroethol Sens Neural Behav Physiol 192, 1327–34.

Caplan, J. S., Williams, A. H. and Marder, E. (2014). Many parameter sets in a multicompartment model oscillator are robust to temperature perturbations. J Neurosci 34, 4963–75.

Chabot, C. C. and Watson, W. H. (2010a). Circatidal rhythms of locomotion in the American horseshoe crab *Limulus polyphemus:* Underlying mechanisms and cues that influence them. Current Zoology 56, 499–517.

Chabot, C. C. and Watson, W. H. (2010b). Horseshoe crab behavior: Patterns and processes. Current Zoology 56, I–Iii.

Chen, R., Jiang, X., Conaway, M. C., Mohtashemi, I., Hui, L., Viner, R. and Li, L. (2010). Mass spectral analysis of neuropeptide expression and distribution in the nervous system of the lobster *Homarus americanus*. J Proteome Res 9, 818–32.

Christie, A. E., Cashman, C. R., Stevens, J. S., Smith, C. M., Beale, K. M., Stemmler, E. A., Greenwood, S. J., Towle, D. W. and Dickinson, P. S. (2008). Identification and cardiotropic actions of brain/gut-derived tachykinin-related peptides (TRPs) from the American lobster Homarus americanus. Peptides 29, 1909–18.

Christie, A. E., Skiebe, P. and Marder, E. (1995). Matrix of neuromodulators in neurosecretory structures of the crab Cancer borealis. J Exp Biol 198, 2431–9.

Clemens, S., Combes, D., Meyrand, P. and Simmers, J. (1998a). Long-term expression of two interacting motor pattern-generating networks in the stomatogastric system of freely behaving lobster. J. Neurophysiol. 79, 1396–408.

Clemens, S., Massabuau, J.-C., Meyrand, P. and Simmers, J. (1999). Changes in motor network expression related to moulting behaviour in lobster: role of moult-induced deep hypoxia. J. Exp. Biol. 202, 817–827.

Clemens, S., Massabuau, J. C., Legeay, A., Meyrand, P. and Simmers, J. (1998b). *In vivo* modulation of interacting central pattern generators in lobster stomatogastric ganglion: influence of feeding and partial pressure of oxygen. J. Neurosci. 18, 2788–99.

Clemens, S., Massabuau, J. C., Meyrand, P. and Simmers, J. (2001). A modulatory role for oxygen in shaping rhythmic motor output patterns of neuronal networks. Respir Physiol 128, 299–315.

Clemens, S., Meyrand, P. and Simmers, J. (1998c). Feeding-induced changes in temporal patterning of muscle activity in the lobster stomatogastric system. Neurosci. Lett. 254, 65–8.

Cooke, I. M. (2002). Reliable, responsive pacemaking and pattern generation with minimal cell numbers: the crustacean cardiac ganglion. Biol Bull 202, 108–36.

Cruz-Bermudez, N. D. and Marder, E. (2007). Multiple modulators act on the cardiac ganglion of the crab, *Cancer borealis*. J Exp Biol 210, 2873–84.

De La Iglesia, H. O. and Hsu Y., W. A. (2010). Biological clocks and rhythms in intertidal crustaceans. Frontiers in Bioscience—Elite. 2010 2, 1394–1404.

DeKeyser, S. S., Kutz-Naber, K. K., Schmidt, J. J., Barrett-Wilt, G. A. and Li, L. (2007). Imaging mass spectrometry of neuropeptides in decapod crustacean neuronal tissues. J Proteome Res 6, 1782–91.

Depledge, M. H. (1983). Photoplethysmography – A non-invasive technique for monitoring heart beat and ventilation rate in Decapod Crustaceans. Comp. Biochem Physiol 77A, 369–371.

Dickinson, E. S., Johnson, A. S., Ellers, O. and Dickinson, P. S. (2016a). Forces generated during stretch in the heart of the lobster *Homarus americanus* are anisotropic and are altered by neuromodulators. J Exp Biol 219, 1187–202.

Dickinson, P. S., Calkins, A. and Stevens, J. S. (2015a). Related neuropeptides use different balances of unitary mechanisms to modulate the cardiac neuromuscular system in the American lobster, *Homarus americanus*. J Neurophysiol 113, 856–70.

Dickinson, P. S., Qu, X. and Stanhope, M. E. (2016b). Neuropeptide modulation of pattern-generating systems in crustaceans: comparative studies and approaches. Curr Opin Neurobiol 41, 149–157.

Dickinson, P. S., Sreekrishnan, A., Kwiatkowski, M. A. and Christie, A. E. (2015b). Distinct or shared actions of peptide family isoforms: I. Peptide-specific actions of pyrokinins in the lobster cardiac neuromuscular system. J Exp Biol 218, 2892–904.

Hamood, A. W., Haddad, S. A., Otopalik, A. G., Rosenbaum, P. and Marder, E. (2015). Quantitative reevaluation of the effects of short- and long-term removal of descending modulatory inputs on the pyloric rhythm of the crab, *Cancer borealis*. ENEURO 2, 0058–14.

Hamood, A. W. and Marder, E. (2014). Animal-to-Animal Variability in Neuromodulation and Circuit Function. Cold Spring Harb Symp Quant Biol 79, 21–8.

Hamood, A. W. and Marder, E. (2015). Consequences of acute and long-term removal of neuromodulatory input on the episodic gastric rhythm of the crab *Cancer borealis*. J Neurophysiol 114, 1677–92.

Heinzel, H. G. (1988). Gastric mill activity in the lobster. I. Spontaneous modes of chewing. J. Neurophysiol. 59, 528–550.

Heinzel, H. G. and Selverston, A. I. (1988). Gastric mill activity in the lobster. III. Effects of proctolin on the isolated central pattern generator. J. Neurophysiol. 59, 566–585.

Hooper, S. L. and Marder, E. (1987). Modulation of the lobster pyloric rhythm by the peptide proctolin. J. Neurosci. 7, 2097–2112.

Hui, L., Xiang, F., Zhang, Y. and Li, L. (2012). Mass spectrometric elucidation of the neuropeptidome of a crustacean neuroendocrine organ. Peptides 36, 230–9.

Johnson, B. and Hooper, S. (1992). Overview of the stomatogastric nervous system. In Dynamic Biological Networks, eds. R. Harris-Warrick E. Marder A. Selverston and M. Moulins), pp. 1–30. Cambridge, MA: MIT Press.

Marder, E. (2012). Neuromodulation of neuronal circuits: back to the future. Neuron 76, 1–11.

Marder, E. and Bucher, D. (2007). Understanding circuit dynamics using the stomatogastric nervous system of lobsters and crabs. Annu Rev Physiol 69, 291–316.

Marder, E., Haddad, S. A., Goeritz, M. L., Rosenbaum, P. and Kispersky, T. (2015). How can motor systems retain performance over a wide temperature range? Lessons from the crustacean stomatogastric nervous system. J Comp Physiol A Neuroethol Sens Neural Behav Physiol 201, 851–6.

Maynard, D. M. (1972). Simpler networks. Ann. NY Acad. Sci. 193, 59–72.

Maynard, D. M. and Dando, M. R. (1974). The structure of the stomatogastric neuromuscular system in *Callinectes sapidus, Homarus americanus* and *Panulirus argus* (decapoda crustacea). Philos. Trans. R. Soc. Lond. (Biol.) 268, 161–220.

McGaw, I. J. and McMahon, B. R. (1996). Cardiovascular responses resulting from variation in external salinity in the dungeness crab, Cancer magister. Physiological Zoology 69, 1384–1401.

McMahon, B. R. (1995). The physiology of gas exchange, circulation, ion regulation and nitrogenous excretion: an integrative approach. In Biology of the Lobster Homarus americanus, (ed. J. R. Factor), pp. 497–517. San Diego: Academic Press.

McMahon, B. R. (1999). Intrinsic and extrinsic influences on cardiac rhythms in crustaceans. Comparative Biochemistry and Physiology a-Molecular and Integrative Physiology 124, 539–547.

McMahon, B. R. (2001a). Control of cardiovascular function and its evolution in Crustacea. J Exp Biol 204, 923–32.

McMahon, B. R. (2001b). Respiratory and circulatory compensation to hypoxia in crustaceans. Respir Physiol 128, 349–64.

Mitra, P. P., Pesaran, B. and (1999). Analysis of dynamic brain imaging data. Biophys J. 76, 691–708.

Morris, L. G. and Hooper, S. L. (1997). Muscle response to changing neuronal input in the lobster (*Panulirus interruptus)* stomatogastric system: spike number-versus spike frequency-dependent domains. J. Neurosci. 17, 5956–5971.

Morris, L. G. and Hooper, S. L. (1998). Muscle response to changing neuronal input in the lobster (*Panulirus interruptus)* stomatogastric system: slow muscle properties can transform rhythmic input into tonic output. J. Neurosci. 18, 3433–3442.

Morris, L. G. and Hooper, S. L. (2001). Mechanisms Underlying Stabilization of Temporally Summated Muscle Contractions in the Lobster (*Panulirus)* Pyloric System. J Neurophysiol 85, 254–268.

O’Leary, T. and Marder, E. (2016). Temperature-robust neural function from activity-dependent ion channel regulation. Current Biology 26, 2935–2941.

Rabiner, L. R. (1989). A tutorial on hidden Markov models and selected applications in speech recognition. Proceedings of the IEEE 77, 257–286 77, 257–286.

Rezer, E. and Moulins, M. (1983). Expression of the crustacean pyloric pattern generator in the intact animal. J. Comp. Physiol. A 153, 17–28.

Rinberg, A., Taylor, A. L. and Marder, E. (2013). The effects of temperature on the stability of a neuronal oscillator. PLoS Comput Biol 9, e1002857.

Robertson, R. M. and Money, T. G. (2012). Temperature and neuronal circuit function: compensation, tuning and tolerance. Curr Opin Neurobiol 22, 724–34.

Soofi, W., Goeritz, M. L., Kispersky, T. J., Prinz, A. A., Marder, E. and Stein, W. (2014). Phase maintenance in a rhythmic motor pattern during temperature changes *in vivo*. J Neurophysiol 111, 2603–2613.

Sullivan, R. E. and Miller, M. W. (1984). Dual effects of proctolin on the rhythmic burst activity of the cardiac ganglion. J Neurobiol 15, 173–96.

Tang, L. S., Goeritz, M. L., Caplan, J. S., Taylor, A. L., Fisek, M. and Marder, E. (2010). Precise temperature compensation of phase in a rhythmic motor pattern. PLoS Biol 8, e1000469.

Tang, L. S., Taylor, A. L., Rinberg, A. and Marder, E. (2012). Robustness of a rhythmic circuit to short- and long-term temperature changes. J Neurosci 32, 10075–85.

Wilkens, J. L. and McMahon, B. R. (1994). Cardiac performance in semi-isolated heart of the crab Carcinus maenas. Am J Physiol 266, R781–9.

Williams, A. H., Calkins, A., O’Leary, T., Symonds, R., Marder, E. and Dickinson, P. S. (2013). The neuromuscular transform of the lobster cardiac system explains the opposing effects of a neuromodulator on muscle output. J Neurosci 33, 16565–75.

Worden, M. K., Clark, C. M., Conaway, M. and Qadri, S. A. (2006). Temperature dependence of cardiac performance in the lobster *Homarus americanus*. J Exp Biol 209, 1024–34.

Zielinski, S. and Portner, H. O. (1996). Energy metabolism and ATP free-energy change of the intertidal worm Sipunculus nudus below a critical temperature. Journal of Comparative Physiology B-Biochemical Systemic and Environmental Physiology 166, 492–500.

